# Interpersonal Synchronization of Brain and Body Tracks Attention and Listening Engagement

**DOI:** 10.64898/2026.08.06.743268

**Authors:** Lotte Lambrechts, Bernd Accou, Jonas Vanthornhout, Bart Boets, Tom Francart

**Author notes:** Corresponding author: Lotte Lambrechts.

## Abstract

**Purpose:** Speech perception is a fundamental part of everyday communication that relies on more than simple identification of words and sentences. Attention and listening engagement both contribute to speech perception, while representing distinct aspects of the listening experience. Attention is typically associated with cognitive focus, whereas listening engagement additionally involves cognitive and affective immersion in sound. Despite their importance, these states remain difficult to disentangle, behaviorally and physiologically. Both have been linked to interpersonal synchronization (the synchronization of biobehavioral signals across individuals), raising questions about what this synchronization actually reflects.

**Method:** In this study, we disentangled attention and listening engagement by independently manipulating both factors within a single experiment. Thirty participants listened to two simultaneously presented streams of meaningful speech and were instructed to focus on only one. Both attended and unattended stimuli were designed to be either engaging or non-engaging. Neural activity was recorded using EEG, while physiological responses were measured using heart rate and electrodermal activity.

**Results:** Interpersonal synchronization was computed from neural and bodily signals, alongside a self-report measure of listening engagement and auditory attention decoding (AAD), a neural measure of selective attention. Interpersonal synchronization of all three modalities significantly predicted listening engagement, whereas neural interpersonal synchronization was the only measure that significantly predicted attention. These findings suggest that attention is primarily driven by cognitive processes represented in the brain, while listening engagement additionally involves affective processes that are more strongly reflected in bodily responses.

**Conclusions:** Overall, this study demonstrates that different forms of interpersonal synchronization reflect distinct dimensions of the listening experience and supports interpersonal synchronization as a potential objective marker of listening engagement.

## Introduction

Every day listening and communication go beyond isolated speech perception (Rönnberg et al., 2013). This becomes, for instance, clear in audiology, where current clinical measures of hearing, which focus on sound identification and speech understanding, often fail to reflect patients’ real-world listening experiences (Carlile et al., 2025; Dornhoffer et al., 2020; Tang et al., 2025). Increasingly, researchers argue that successful listening depends on higher-order listening states, like attention and listening engagement (Herrmann & Johnsrude, 2020; Pichora-Fuller et al., 2016). Understanding how these listening states contribute to speech perception, and how they can be objectively measured, remains an important challenge.

Speech perception involves transforming acoustic input into meaningful linguistic representations through consecutive auditory and linguistic processing (Friederici, 2011; Hickok & Poeppel, 2007). Beyond these processes, successful speech comprehension also relies on cognitive mechanisms such as working memory and executive functioning (Kaya & Elhilali, 2017; Pichora-Fuller et al., 2016; Rönnberg et al., 2013). Over the past decade, researchers have worked to model how these processes unfold in the brain (Brodbeck & Simon, 2020; Crosse et al., 2021; Vanthornhout et al., 2018). The brain tracks heard speech by synchronizing with features of the acoustic speech signal, a phenomenon known as neural tracking (Gillis et al., 2022). Neural tracking can be quantified by training models, such as linear decoders, that relate neural responses recorded via EEG, from people listening to speech, to the speech signal (Crosse et al., 2021; Gillis et al., 2022). Neural tracking to speech is commonly considered a reliable proxy for speech understanding (Iotzov & Parra, 2019; Vanthornhout et al., 2018). As a result, it has become an important tool in both auditory neuroscience and clinical audiology (Brodbeck & Simon, 2020; Crosse et al., 2021; Gillis et al., 2022; Iotzov & Parra, 2019; Van Hirtum et al., 2023). However, neural tracking does not solely reflect speech perception, but is strongly affected by attention to speech (Iotzov & Parra, 2019; Vanthornhout et al., 2019). Attention to speech refers to the cognitive process of selectively directing and sustaining focus on spoken language while filtering out irrelevant information (Kaya & Elhilali, 2017; Pichora-Fuller et al., 2016). It also plays an essential role in the retention of spoken information (Song et al., 2021). Previous research has demonstrated that neural tracking significantly decreased when participants were distracted from the speech stimulus (e.g., by watching an incongruent movie). Building on this idea, neural tracking measures can be applied in more complex listening situations, such as those involving multiple speakers, requiring people to focus on one speaker while ignoring others (O’Sullivan et al., 2015). To assess how well a person can attend to one speaker and ignore another, auditory attention decoding (AAD) can be employed (Das et al., 2020; Ding & Simon, 2012; Mesgarani & Chang, 2012). Here, neural tracking of both the attended and ignored speaker is compared to determine which one the person is focusing on. For the attended speaker, neural tracking will be higher than for the ignored one. In general, the AAD method contributes to our understanding of speech perception and its modulation by attention (Geirnaert et al., 2021; Gillis et al., 2022) and is already clinically implemented to improve hearing aid performance by more accurately identifying and, consequently, selectively amplifying the desired speaker (Geirnaert et al., 2021; Roebben et al., 2024).

In addition, the literature has presented interpersonal synchronization as an alternative measure of attention (Dmochowski et al., 2012; Madsen & Parra, 2025; Stuldreher et al., 2020). This concept refers to the alignment of brain and bodily responses across individuals during a shared experience.

In this context, listening is regarded as a shared experience, and interpersonal synchronization of biobehavioral signals has been shown to emerge when different people listen to the same sound stream, even in the absence of direct interaction (Madsen & Parra, 2022, 2024; Pérez et al., 2021). Madsen & Parra (2022, 2024) demonstrated this by showing that individuals listening to the same speech material exhibit increased interpersonal synchronization compared to conditions without stimulation, regardless of whether they were listening simultaneously or at different times or in different spaces. They further showed that interpersonal synchronization decreased when listeners were distracted. Despite these seemingly clear associations, recent literature suggests that interpersonal synchronization may reflect not only attention, but also listening engagement, raising questions about what the measure truly captures.

Listening engagement is increasingly recognized as an important factor in speech perception. Theories of narrative engagement proposed by Busselle & Bilandzic (2009), together with the Model of Listening Engagement developed by Herrmann & Johnsrude (2020), provide the foundation for current conceptualizations of this construct. Listening engagement refers to the degree to which listeners become immersed in an auditory experience (Busselle & Bilandzic, 2009). This experience involves both a cognitive component, reflected in increased cognitive and integrative processing (Herrmann & Johnsrude, 2020), and an affective component associated with emotional involvement and empathy (Busselle & Bilandzic, 2009). Although the models by Herrmann & Johnsrude (2020) and Busselle & Bilandzic (2009) approach the concept from different perspectives, they share several commonalities. Both models characterize engagement as a flow-like state, marked by reduced self-awareness during listening (Cziksentmihalyi, 1990). This state emerges when listeners are motivated to engage by anticipated/ perceived enjoyment, curiosity, or a desire to acquire knowledge (Bradbury, 2023; Herrmann & Johnsrude, 2020; Liang et al., 2018). This contrasts with attention, which primarily refers to the conscious focus directed to speech stimuli (Pichora-Fuller et al., 2016). Based on this literature, we hypothesize that attention may function as a prerequisite for listening engagement, although engagement itself may also influence how attention is sustained and allocated during listening (Bradbury, 2023). In summary, listening engagement extends beyond attention by incorporating both sustained cognitive processing and affective involvement in the listening experience.

Currently, listening engagement is evaluated through validated self-report scales in which people rate their engagement by responding to statements on a Likert scale (e.g., the narrative engagement scale (NES) (Busselle & Bilandzic, 2009), the transportation scale (Green & Brock, 2000), the narrative absorption scale (Kuijpers et al., 2014)). Despite their usefulness, these scales have inherent limitations, such as response bias and the need for active reflection by the respondent. Following these limitations, the need for a new approach to measuring listening engagement was large. Some recent papers have suggested that interpersonal synchronization between brain and body signals may serve as a measure of listening engagement (Chatterjee et al., 2021; Czepiel et al., 2025; Irsik et al., 2022). In our own lab, we were the first to directly manipulate listening engagement and show that interpersonal synchronization increased significantly with increased listening engagement (Lambrechts et al., 2025). However, this raises important conceptual questions, leaving it unclear whether this interpersonal synchronization measure mainly reflects attention, listening engagement, or both.

To disentangle these competing interpretations, we simultaneously manipulated attention and listening engagement while combining two key measurement approaches in auditory neuroscience: auditory attention decoding (AAD) and interpersonal synchronization. We expected both AAD and interpersonal synchronization to reflect differences arising from our experimental manipulations (Research question 1). We hypothesized that interpersonal synchronization would capture variance associated with listening engagement beyond that explained by attention alone. Accordingly, we expected attention and listening engagement to emerge as distinct yet related listening states, each contributing uniquely to interpersonal synchronization (Research question 2). Finally, we sought to further validate interpersonal synchronization as a physiological marker of listening engagement by comparing it to self-report listening engagement measures (Research question 3).

## Methods

In this study, we invited normal-hearing adults to listen to two streams of speech stimuli at the same time. The presented stimuli were either engaging or non-engaging, and they were instructed to attend to only one of them.

### Participants

For this study, 30 participants were recruited. All participants had normal hearing, as identified by pure-tone audiometry (Fletcher Index below 20 dB HL). Additionally, through a brief screening questionnaire, we ensured the absence of any additional disorders or neural or developmental abnormalities. Participants had an average age of 22 years, ranging from 18 to 30 years. 14 of 30 participants were male, resulting in a sex-balanced dataset. Prior to any experimental testing, participants provided informed consent to participate in our study, which was approved by the Medical Ethics Committee UZ Leuven/Research (KU Leuven) with reference S68492.

### Experimental Preparation and Setup

Participants were seated in a soundproof room in a comfortable chair. In front of them, at a distance of 110 cm, a computer monitor was placed, on which they could follow the experimental procedure. Two Genelec (8020a) loudspeakers were placed at a distance of 140 cm and in front of the participant at an angle of ± 25°. See figure 1 for a full overview.

**Figure 1:**
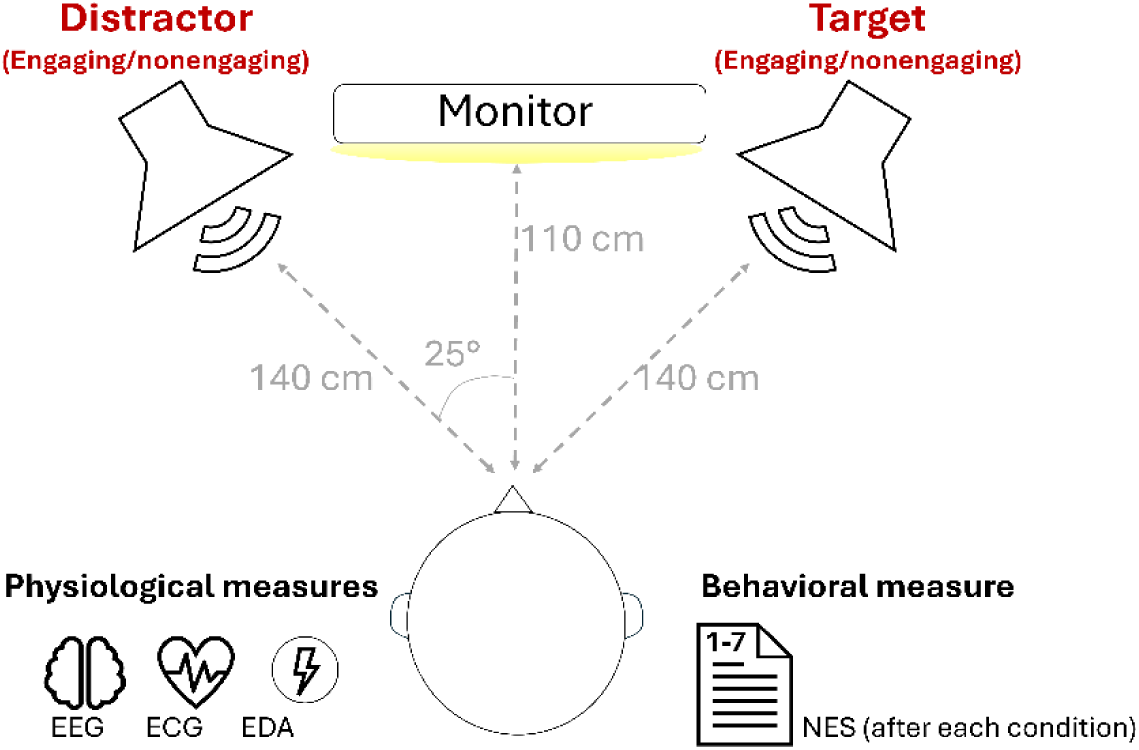
Overview of the experimental setup with positioning of the two loudspeakers and screen in relation to the participant, and the included measures.

Stimuli were presented using APEX stimulation software (Francart et al., 2008). All of them were presented at 60 dB A. To ensure equal loudness, we took some precautions during stimulus preparation. First, we smoothed the root-mean-square (RMS) within each stimulus using MATLAB’s smoothdata function with a 20-ms moving-average window, and then rescaled the RMS across all stimuli to a common level. Prior to the experiments, the full setup with two loudspeakers was calibrated using Brüel & Kjær calibration equipment in dB A (Brüel & Kjær, Naerum, Denmark). As calibration stimuli, we used speech-weighted noise calculated for each included stimulus.

### Experimental test session - procedure

Participants were invited to our hearing lab for a 2-hour experimental session.

- **Preparation**: After a brief explanation of the experiment, participants signed the informed consent form. Next, we administered a brief screening questionnaire to verify inclusion criteria and gather demographic information, and performed pure-tone audiometry to confirm that each participant had normal hearing.
- **Brain & body sensor application**: All brain and body responses were recorded using a BioSemi ActiveTwo system (BioSemi, The Netherlands) at a sample rate of 8192 Hz and later down- sampled. The complete sensor application took about 15 minutes.

- *Electroencephalography (EEG):* Participants were fitted with a 64-channel BioSemi EEG system according to the 10-20 system. We used a conductive gel to ensure there was a good connection between each electrode and the skull. Proper application was evaluated by ensuring that no signals were visually noisy and that the electrode DC offsets (relative to CMS) were within the -25 to 25 mV range.
- *Electrocardiography (ECG)*: We applied two Biosemi flat electrodes to the participants’ bodies. One was placed 2 cm below the left clavicle, and the other in the left lumbar region (lower back). This setup is consistent with previous studies (Lambrechts et al., 2025; Madsen & Parra, 2022, 2024). To ensure proper measurement, we applied conductive gel and used tape to prevent the sensors from moving. Before beginning the measurement, we visually checked whether the heart rate R-peak was identifiable in the signal.
- *Electrodermal activity (EDA):* We applied a BioSemi galvanic skin response sensor with two electrodes placed on the left pointer and middle fingers. We used a conductive gel to ensure proper response acquisition and applied tape to prevent sensor movement.
- **Full experimental procedure**: In all listening conditions, we exposed participants to 2 streams of continuous natural podcast-style, speech stimuli, simultaneously presented via two spatially separated loudspeakers. Participants were instructed to attend only to the conversation presented through the right loudspeaker. The speech stimuli consisted of continuous speech from female speakers to minimize acoustic variability. All stimuli were approximately 15 minutes in duration and were categorized as either engaging or non-engaging. The engaging condition comprised excerpts from the Flemish podcast *De Volksjury*, in which two female hosts discuss crime-related stories in an engaging manner. The non-engaging condition consisted of excerpts from recordings of the Flemish Parliament (2015), in which speakers discussed political topics that were not necessarily temporally or personally relevant to participants. The study included the following conditions, which were defined by different combinations of these stimulus types:

| Condition | Distractor (left loudspeaker) | Target (right loudspeaker) |
| --- | --- | --- |
| **Condition 1** | Non-engaging stimulus<br>*'Vlaams Parlement' part 1* | Engaging stimulus<br>*'De Volksjury – Erikson Tweeling' part 1* |
| **Condition 2** | Engaging stimulus<br>*'De Volksjury – Maddie McCann' part 1* | Engaging stimulus<br>*'De Volksjury – Erikson Tweeling' part 2* |
| **Condition 3** | Engaging stimulus<br>*'De Volksjury – Maddie McCann' part 2* | Non-engaging stimulus<br>*'Vlaams Parlement' part 2* |
| **Condition 4** | Non-engaging stimulus<br>*'Vlaams Parlement' part 4* | Non-engaging stimulus<br>*'Vlaams Parlement' part 3* |

The combinations of specific stimuli were always the same; the order of conditions was randomized across participants to control for learning and/or fatigue effects. We ensured that condition 1 always came before condition 2 to ensure the story was told in chronological order. During the listening task, participants were encouraged to remain seated, calm and comfortable and to maintain their attention on the instructed stimulus to the best of their ability.

After each listening condition, we administered the narrative engagement scale (NES) (Busselle & Bilandzic, 2009), translated to Dutch, to assess perceived listening engagement behaviorally. This validated scale consists of 12 7-point Likert statements and is categorized into 4 subscales (Narrative Understanding, Attention Focus, Narrative Presence and Emotional Engagement). In the analyses, we calculated average ratings for each participant and condition across all statements, yielding 4 values per participant. The full protocol, consisting of 4 listening conditions, each with a duration of 15 minutes, resulted in an overall testing time of ± 1.5 hours, including problem management and breaks.

### Signal processing

All signal processing was done in MATLAB (version 24.2.0.2740171 (R2024b), Natick, Massachusetts: The MathWorks Inc)

### Pre-processing of different brain-body responses and stimuli

#### EEG

EEG data were preprocessed to reduce noise and ensure comparability across subjects. First, a first- order Butterworth high-pass filter at 0.5 Hz was applied to remove signal drift. The data were then resampled using the MATLAB resample function from 8192 to 1024 Hz. Artifact reduction was performed using a multi-channel Wiener filter targeting eyeblinks (Somers et al., 2018). Subsequently, the data were re-referenced using common average referencing to attenuate noise shared across channels. The signal was further downsampled to 64 Hz and band-pass filtered between 0.5 and 8 Hz to extract the delta and theta bands. Finally, all channels were z-scored per trial to facilitate comparisons across subjects and conditions. Silences were not excluded. These preprocessing steps were completed prior to training neural tracking decoders and performing auditory attention decoding (AAD) analyses.

#### Heart rate

ECG processing and extraction of the instantaneous heart rate were performed following the procedure described by Pérez et al (2021), using their publicly available code with minor adaptations. First, we band-pass filtered the ECG between 0.5 and 25 Hz to remove slow drift and preserve relevant cardiac activity, which typically occurs at lower frequencies. Next, we re-referenced the channels to one another, yielding a single, clear ECG signal. We extracted heart rate peaks and computed the instantaneous heart rate by measuring the time intervals between successive peaks (Pérez et al., 2021). Finally, the resulting heart rate time series was resampled to 64 Hz.

#### EDA

Signal processing followed the method described by Marci et al (2007). We smoothed the EDA response by convolving the data with a 1.5 s Bartlett window to reduce noise and ensure accurate response extraction. From this smoothed EDA data, we calculated the derivative by computing differences between consecutive samples and consecutively applied a sigmoid transformation (i.e., y = 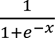, with x = EDA amplitude) to the output. Finally, the resulting processed EDA response was resampled to 64 Hz.

#### STIMULI

All stimuli had a sample rate of 48 kHz. To calculate neural tracking and AAD performance, we extracted the stimulus envelope as a feature for all stimuli. We applied a Gammatone Filterbank, filtering the stimulus into 28 erb-spaced bands spanning 50-5000 Hz. Next, we extracted the slow modulations by taking the absolute value within each band and compressed the signal by raising it to the power of 0.6 (Biesmans et al., 2017). Finally, we calculated the actual stimulus envelope by averaging the 28 subbands. To match our body signals, we downsampled the envelope to 64 Hz. This analysis was applied to all engaging and non-engaging stimuli.

### Calculation of Physiological Outcome Measures

#### Neural tracking

In preparation for the AAD calculation, we calculated neural tracking. This was quantified using a backward-modeling approach in which a linear decoder was trained to reconstruct the speech envelope from the EEG signal (Crosse et al., 2021). We implemented leave-one-fold-out cross-validation, with each test fold lasting 30 seconds and the training data consisting of all other EEG recordings from that condition for each person. The model was trained using ridge regression, with the regularization parameter (λ) set to the maximum absolute value of the EEG covariance matrix to prevent overfitting. We included time lags from 0 to 250 milliseconds. This decoder was trained on the target speech to reconstruct the envelope, which was then correlated with the envelopes of both the target and distractor stimuli, yielding separate neural tracking values for each.

#### Auditory attention decoding

We calculated AAD accuracy, reflecting how often neural tracking of target speech exceeded neural tracking of the distractor. For each fold, the size of the correlation with the attended speech was compared to the size of the unattended one, resulting in ‘1’ if the target speech (instructed to attend) was higher and 0 if the opposite was true. With ∼15-minute stimuli segmented into 30-second folds, this yielded 30 comparisons per participant per condition, which were finally converted into proportions reflecting AAD accuracy. Performance exceeding chance level was evaluated using a binomial test (p = 0.5, 30 folds).

#### Intersubject correlations

To quantify interpersonal synchronization for each modality individually, we calculated intersubject correlations (ISC) for each participant, for each condition. Prior to calculating ISC, for EEG, we applied correlated component analysis (corrCA) (Parra et al., 2018). This method computes spatial filters that, when applied, transform 64-channel EEG data into 64 components that maximize correlation between subjects. The filter weights were estimated on the EEG of a subset of participants (28) and subsequently applied to the EEG data held-out subject pairs, resulting in 2 sets of 64 EEG components. Further EEG-synchronization analyses were conducted only with the first component. For each participant, we Spearman-correlated their preprocessed physiological signals (EEG, heart rate (HR), and EDA) with those of all other participants within the same condition. With 30 participants per condition, this yielded 29 pairwise correlations per participant. They were averaged to obtain a single ISC value per participant per condition, separately for each physiological measure, reflecting how well each person correlated with the entire group. To assess statistical significance, we constructed a null distribution as follows: Each participant’s time series (EEG (corrCA), HR, or EDA) was correlated with those of all other participants in the other conditions, resulting in a null distribution of 87 values (i.e., 3 other conditions x 29 other participants). Next, to approximate normality, we Fisher-transformed both the observed correlation value and the null distribution. Because the observed ISC values were obtained by averaging 29 correlation coefficients, we adjusted the null distribution to ensure a fair comparison with comparable variation. To do this, we transformed the standard deviation by applying: 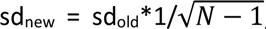, where N = 30. The p-value was calculated by subtracting one from the cumulative distribution function with zero mean and SD, calculated as before, from -infinity to the Fisher-transformed correlation.

#### Statistical analysis

All statistical analyses were performed in R (R version 4.3.1 (2023-06-16), using ‘ggplot2’ for plotting and ‘nlme’ for calculating regression models. For all analyses, we applied regression analysis. The continuous variables of interest are all bounded, which can undermine the reliability and interpretability of a linear mixed-effects model (Akram et al., 2023; Hutmacher et al., 2011). To address this, we use non-linear mixed-effects models to capture a linear yet bounded relationship, where all variables are interpreted as bounded rather than infinite. The included outcome measures and their corresponding bounds are shown in Table 1.

**Table 1:** Outcome measures and their included bounds.

| Outcome measures/ Predictors | Bounds |
| --- | --- |
| Narrative listening engagement (subjective ratings) | 1 – 7 |
| Brain synchronization | -1 – 1 |
| HR synchronization | -1 – 1 |
| EDA synchronization | -1 – 1 |
| AAD accuracies | 0 – 1 |

The included categorial variables, target and distractor, both had two levels: engaging and non-engaging. To treat them as predictors in the NLME models, we used effect coding, with ‘engaging’ coded as 1 and ‘non-engaging’ coded as -1.

To answer **research question 1**, we used 5 regression models to predict the effects of listening engagement induced by both target and distractor speech on the outcome measures. The NLME models included ‘target’ and ‘distractor’ as fixed effects, along with their interaction. Participants were included as a random effect. The specific models we evaluated are shown in Table 2.

**Table 2:** Outcome measures and their included bounds.

| RQ | Models used to answer each sub-question of Research Question 1 |
| --- | --- |
| 1a | *Behavioral responses (1 - 7)* $\sim target + distractor + target * distractor + (1 \vert participant)$ |
| 1b | *Brain synchronization (-1 - 1)* $\sim target + distractor + target * distractor + (1 \vert participant)$ |
| 1b | *HR synchronization (-1 - 1)* $\sim target + distractor + target * distractor + (1 \vert participants)$ |
| 1b | *EDA synchronization (-1 - 1)* $\sim target + distractor + target * distractor + (1 \vert participant)$ |
| 1c | *AAD accuracy (0 - 1)* $\sim target + distractor + target * distractor + (1 \vert participant)$ |

To address **research question 2,** we modeled the relative contributions of attention and listening engagement to interpersonal synchronization. We used non-linear mixed-effect models with attention (AAD accuracies) and listening engagement (subjective ratings) as predictors of EEG, HR and EDA synchronization (bounded between -1 and 1). The general model structure was:

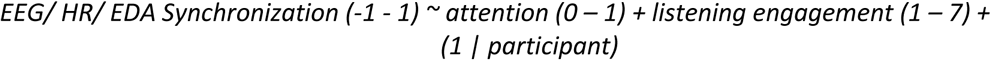

For **research question 3**, we modeled the relationship between behavioral and physiological listening engagement while controlling for experimental conditions. To do so, we employed NLME models that included both interpersonal synchronization and target and distractor conditions as fixed effects, along with their interactions with physiological interpersonal synchronization. Including the interaction allowed us to test whether the association between behavioral engagement and physiological synchronization differed across listening conditions.

The general model structure was:

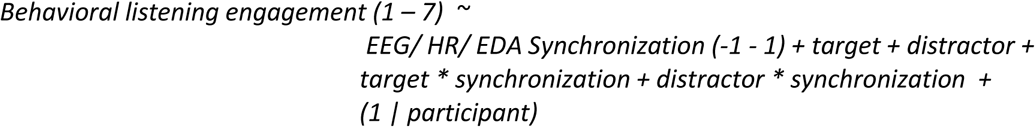

### Permutation testing

To calculate p-values of each estimate in the NLME models, we applied permutation testing. For each model, for each main and interaction effect, we randomized the labels a thousand times for the target, the distractor and both simultaneously (to test the interaction) within each subject. From this, we generated null distributions containing randomly calculated estimates for each effect for each model. Finally, we calculated a two-tailed p-value as the proportion of null values with absolute values at least as large as the observed correlation. By evaluating each predictor relative to its own null distribution, we ensured that their effects could be compared, enabling a reliable assessment of their relative contributions to each outcome variable.

## Results

### RQ1 - a: Effect of listening engagement on the Narrative Engagement Scale

Participants reported self-perceived listening engagement by rating 12 NES statements on a 7-point Likert scale. They were instructed to evaluate only the stimulus they were instructed to attend to. As shown in *Figure 2*, participants reported higher listening engagement when the target stimulus was engaging (blue data). A regression analysis using a non-linear mixed-effects model revealed significant main effects of target speech and, to a smaller extent, distractor speech. No significant interaction effect between target and distractor engagement was observed. These results suggest that both attending to an engaging stimulus and ignoring an engaging distractor independently increased reported listening engagement. Full model output is presented in Table 3.

**Figure 2:**
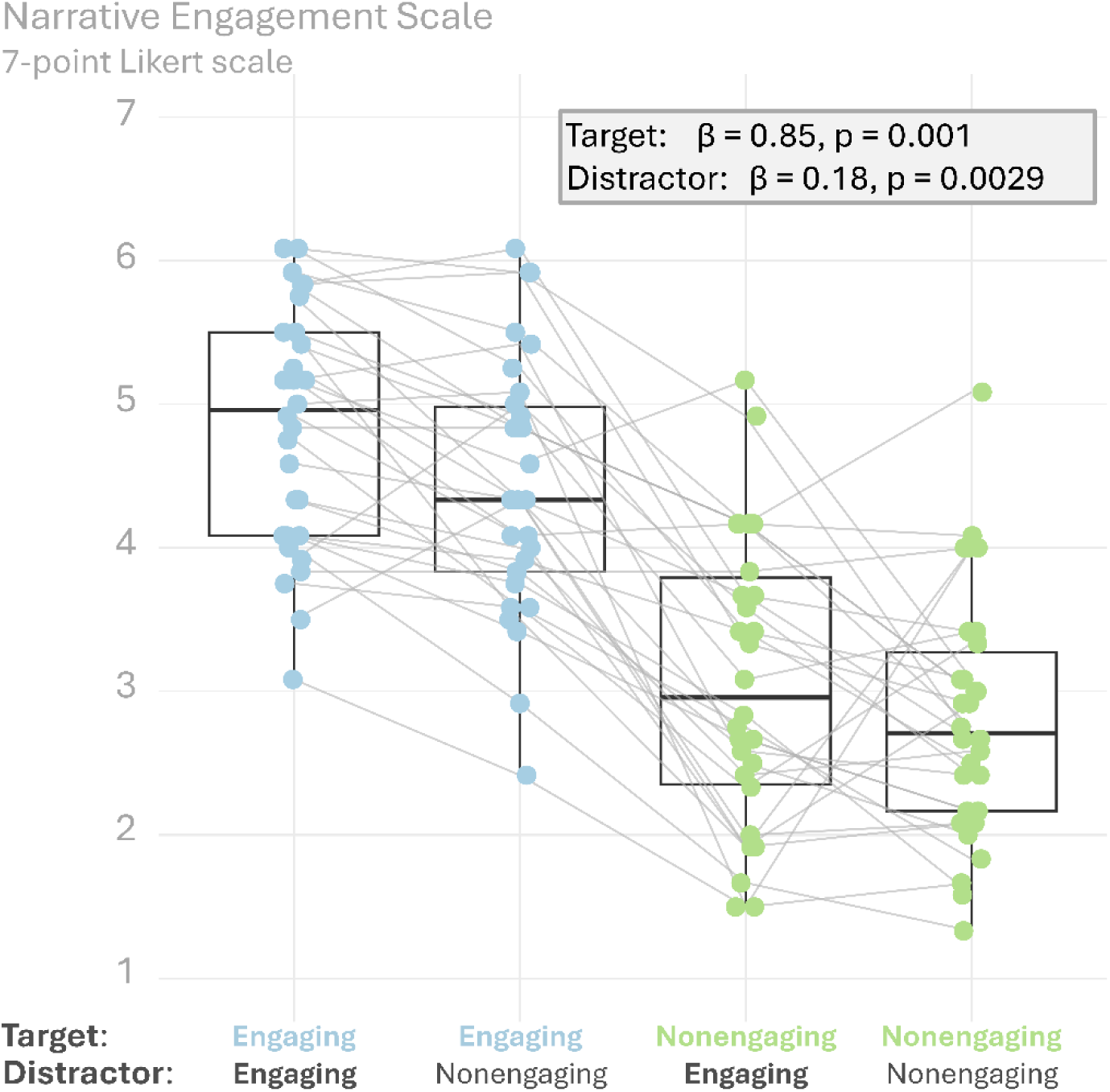
Participants’ subjective listening engagement ratings of the target stimulus. Subjective ratings were significantly higher for engaging target stimuli than for non-engaging target stimuli. Additionally, listening engagement scores for the target stimulus were also higher when the distractor stimulus was engaging instead of non-engaging. Each datapoint is the average score across these 12 statements of the NES for each condition and person.

**Table 3:** Non-linear mixed effects models for behavioral listening engagement.

| *Behavioral responses ~ target + distractor + target \* distractor + (1 \| participants)* |  |  |  |  |
| --- | --- | --- | --- | --- |
|  | Estimate | Std Err | df | p-value |
| *Intercept* | 3.79 | 0.13 | 87 |  |
| *Target* | 0.85 | 0.06 | 87 | 0.001 |
| *Distractor* | 0.18 | 0.06 | 87 | 0.0029 |
| *Target \* Distractor* | 0.044 | 0.06 | 87 | 0.63 |

### RQ1 - b: Effect of listening engagement on interpersonal synchronization

Figure 3 illustrates the effects of listening engagement (engaging vs. non-engaging) on interpersonal synchronization across the three physiological modalities (brain, heart rate, and electrodermal activity) for both the target and the distractor speech. Significant synchronization could be found for all modalities, but not for all participants/ conditions (see Figure 3, grey dots). Across all modalities, interpersonal synchronization was higher when participants listened to an engaging target stimulus compared to a non-engaging one. The NLME models confirmed a significant main effect of the target across all modalities: brain synchronization, HR synchronization, and EDA synchronization and a significant main effect of distractor for EDA synchronization. No further significant main effect of the distractor nor any interaction effect could be identified, meaning interpersonal synchronization is predominantly influenced by what is attended and not by what is ignored. Full results are presented in Table 4.

**Figure 3:**
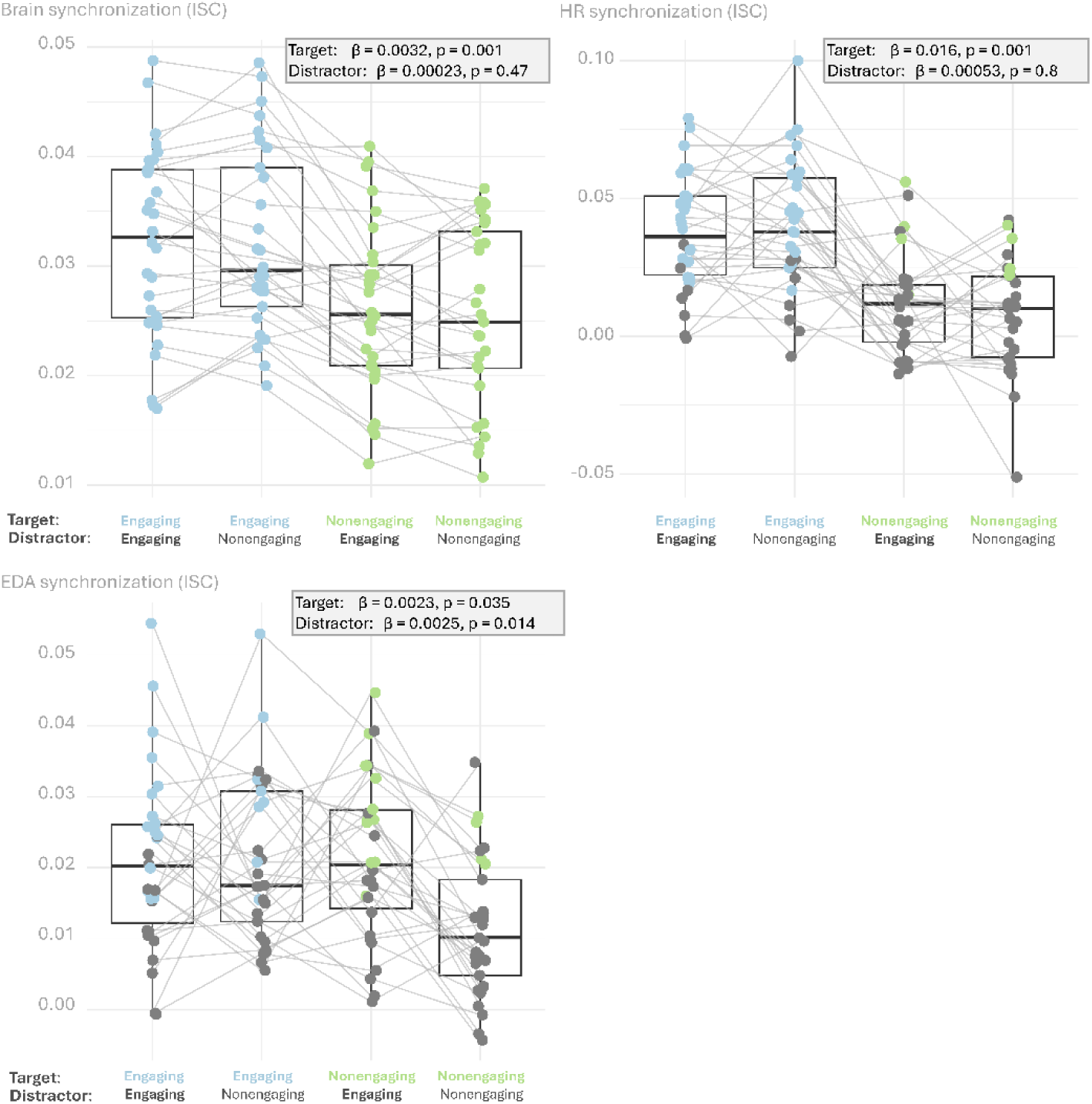
Interpersonal synchronization of neural, HR and EDA responses is significantly influenced by listening engagement in the target speech, but not by listening engagement in the distractor speech. Each datapoint represents the average correlation of one participant in a condition, correlated with all others in that condition. Grey dots indicate insignificant interpersonal synchronization values and were obtained by comparing each correlation to its own null distribution (see methods).

**Table 4:**
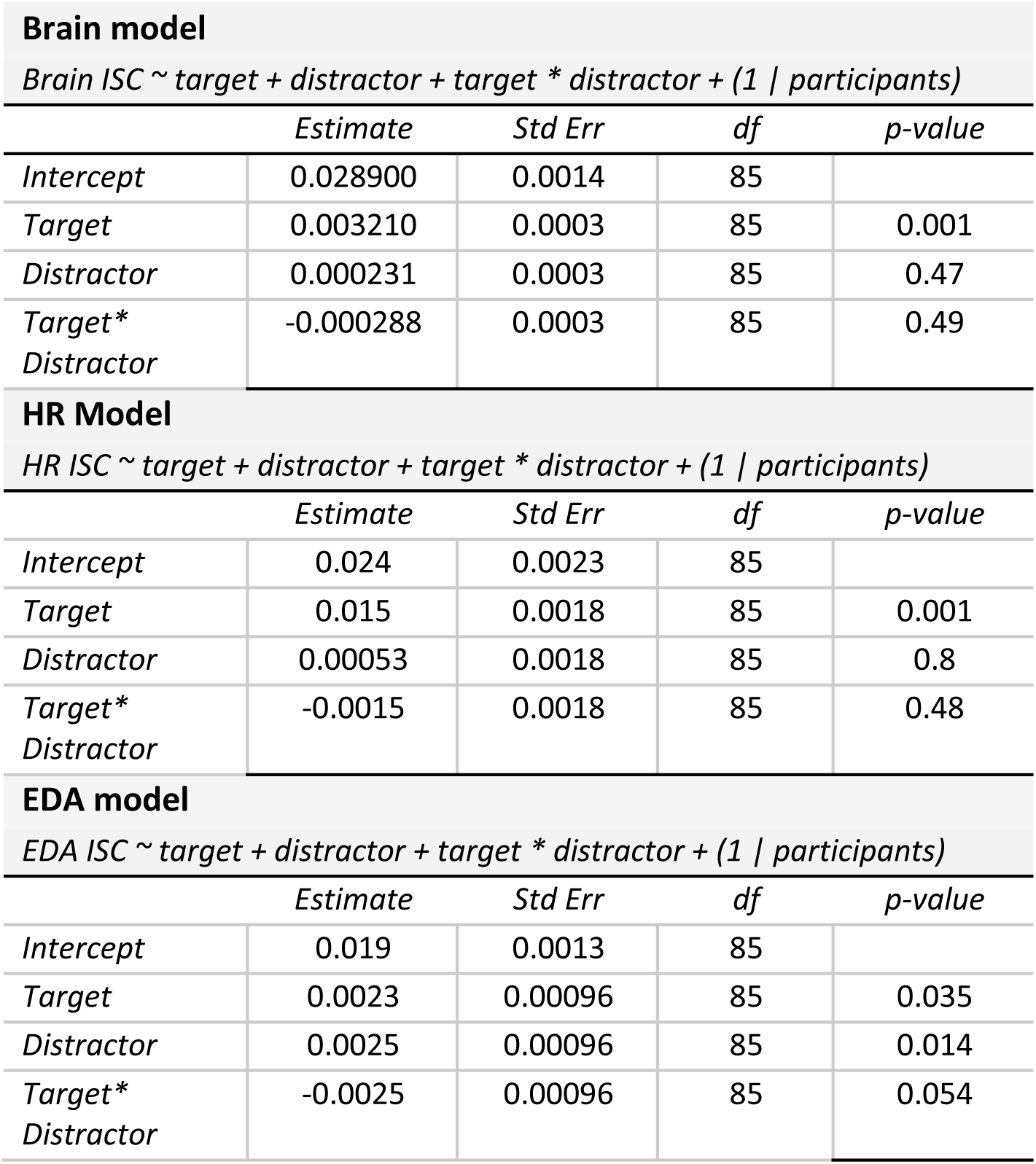
Results from the non-linear Mixed Effects models for the synchronization values.

### RQ1 - c: Effect of engagement on AAD accuracies

Figure 4 illustrates the influence of stimulus engagement on auditory attention decoding performance, as measured by AAD accuracy. Significant AAD accuracies were observed across all experimental conditions, although not every participant achieved significant AAD within each condition (see grey dots in Figure 4). Overall, AAD accuracies were higher when the distractor speech was non-engaging compared to when it was engaging and to a smaller extent when the target was engaging, compared to non-engaging. Statistical analyses revealed significant main effects of both target engagement and distractor engagement on AAD accuracy. However, no significant interaction effect between target and distractor engagement was found. These findings indicate that AAD performance is sensitive to the level of engagement in both target and distractor speech streams. Full results can be found in table 5.

**Figure 4:**
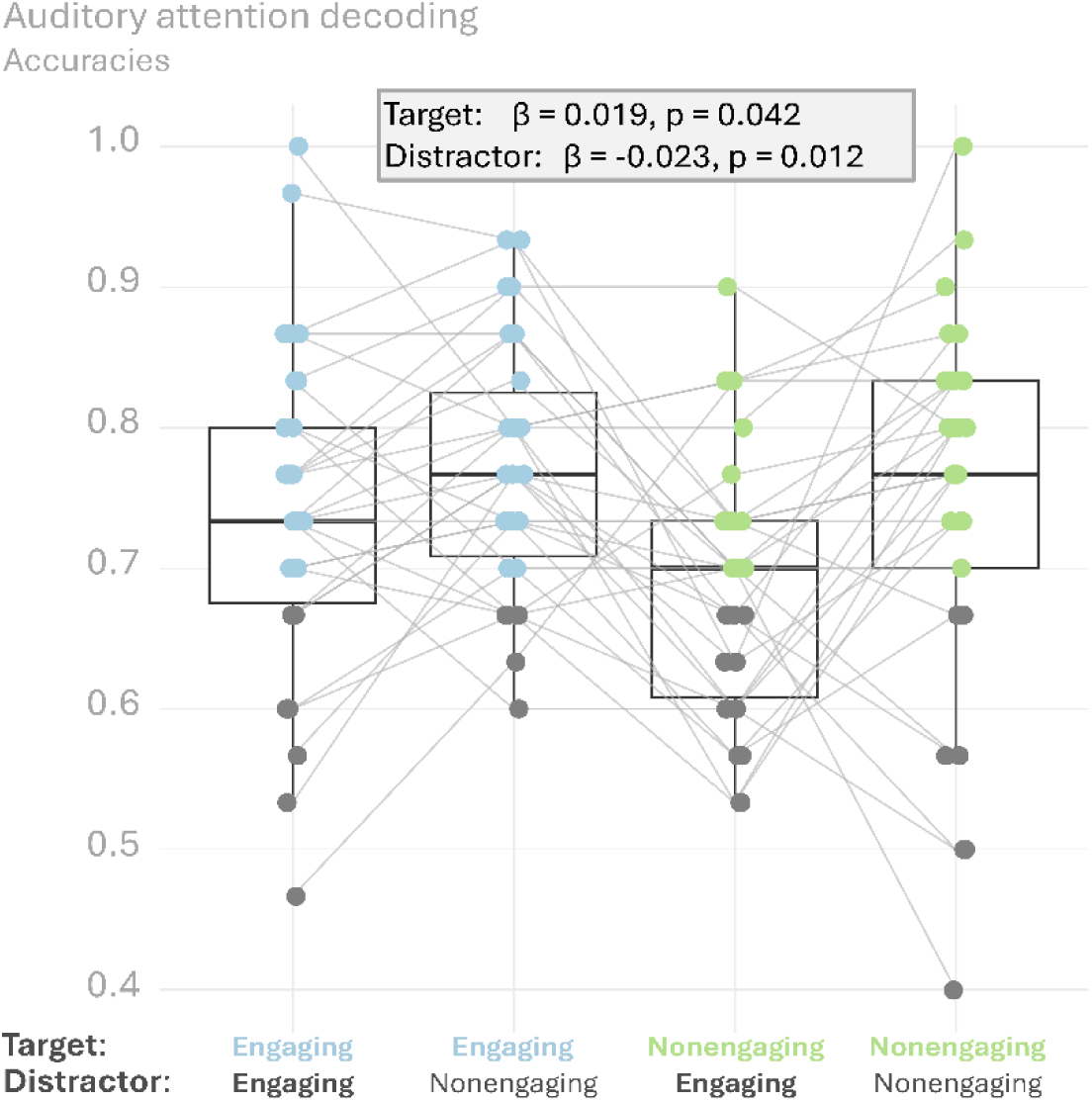
AAD accuracies were significantly higher when target speech was engaging in comparison to when it was non-engaging, and when distractor speech was non-engaging in comparison to when it was engaging. Grey dots indicate significance for the AAD accuracies, identified by binomial testing.

**Table 5:** Results from the non-linear Mixed Effects models for AAD accuracies.

| AAD accuracies |  |  |  |  |
| --- | --- | --- | --- | --- |
| AAD accuracies ∼ target + distractor + target * distractor + (1 \| participants) | | | | |
|  | Estimate | Std Err | df | p-value |
| Intercept | 0.74 | 0.014 | 86 |  |
| Target | 0.019 | 0.008 | 86 | 0.042 |
| Distractor | -0.023 | 0.008 | 86 | 0.012 |
| Target*<br>Distractor | 0.0064 | 0.008 | 86 | 0.45 |

### RQ2: Predictive value of engagement and attention for synchronization

For research question 2, we predicted the relative contributions of attention (AAD accuracies) and listening engagement (behaviorally scored - NES) to interpersonal synchronization across modalities using an NLME model. *Figure 5* contains scatter plots showing this relationship. Overall, the pattern is consistent: interpersonal synchronization increased both with behavioral listening engagement and with focused attention (as measured by AAD accuracies). This pattern is confirmed by the NLME models. Behavioral engagement is a significant predictor of synchronization across all three modalities. Attention was also identified as a significant predictor, but only for brain synchronization and not for HR or EDA synchronization. We used permutation testing to assess the significance and relative importance of each individual predictor. Across all modalities, we found that behavioral listening engagement had a stronger association with synchronization (lower p-values) than attention, even though synchronization estimates were smaller. The full results from these models are presented in Table 6.

**Figure 5:**
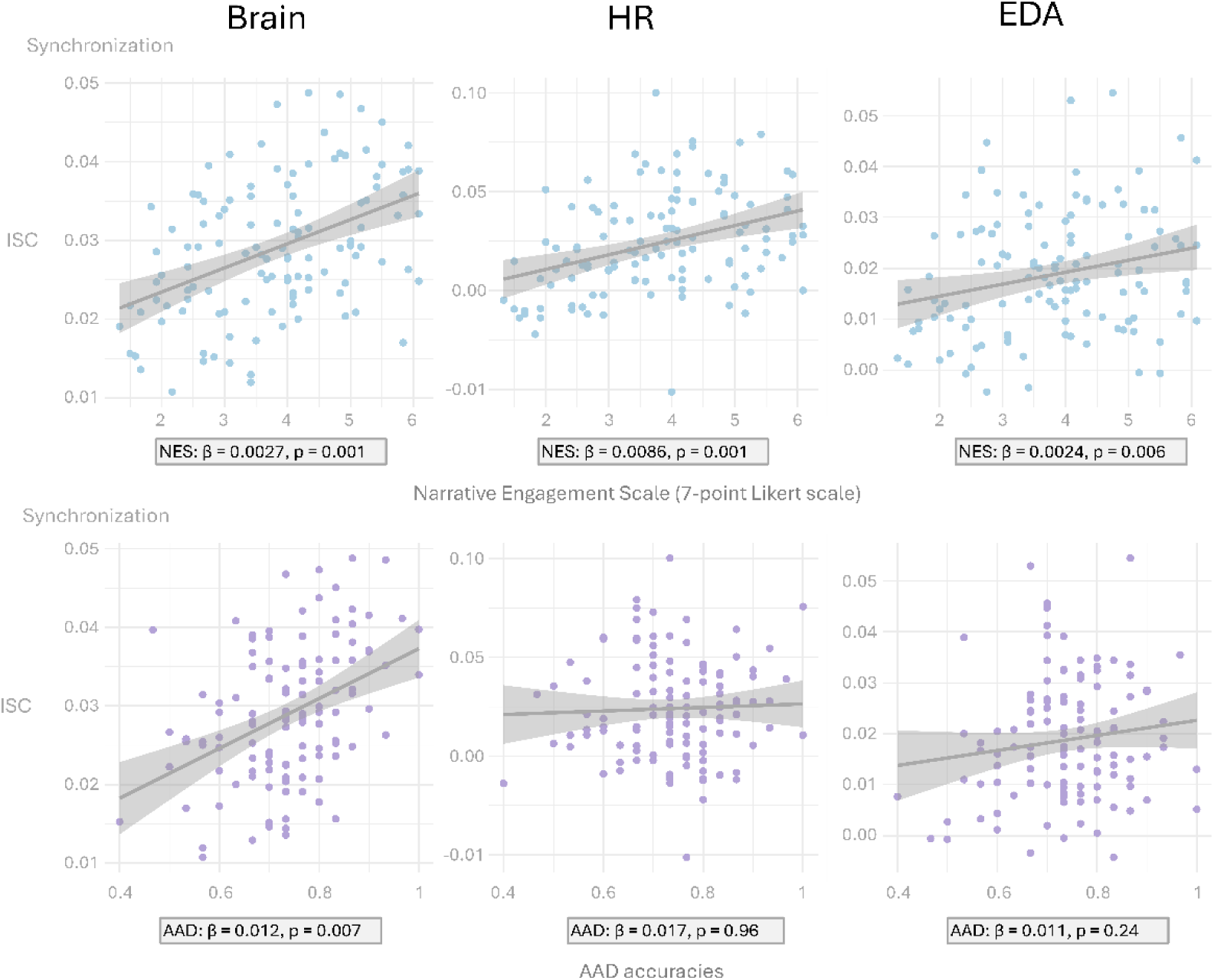
Top row: Self-rated listening engagement is significantly related to neural, HR and EDA interpersonal synchronization when controlled for attention (AAD accuracies). Bottom row: Attention (i.e., AAD accuracies), controlled for listening engagement, also significantly relates to neural interpersonal synchronization but not to HR and EDA synchronization.

**Table 6:** Non-linear mixed effects models reporting the relation between synchronization, attention and engagement.

| **Brain model** |  |  |  |  |
| --- | --- | --- | --- | --- |
| *EEG ISC ~ attention + listening engagement + (1 \| participants)* |  |  |  |  |
|  | *Estimate* | *Std Err* | *df* | *p-value* |
| *Intercept* | 0.01000 | 0.0033 | 86 |  |
| *Engagement* | 0.0027 | 0.00032 | 86 | 0.001 |
| *Attention* | 0.012 | 0.0039 | 86 | 0.007 |
| **HR Model** |  |  |  |  |
| *HR ISC ~ attention + listening engagement + (1 \| participants)* |  |  |  |  |
|  | *Estimate* | *Std Err* | *df* | *p-value* |
| *Intercept* | -0.021 | 0.016 | 86 |  |
| *Engagement* | 0.0086 | 0.0018 | 86 | 0.001 |
| *Attention* | 0.017 | 0.02 | 86 | 0.96 |
| **EDA model** |  |  |  |  |
| *EDA ISC ~ attention + listening engagement + (1 \| participants)* |  |  |  |  |
|  | *Estimate* | *Std Err* | *df* | *p-value* |
| *Intercept* | 0.0019 | 0.0077 | 86 |  |
| *Engagement* | 0.0024 | 0.00087 | 86 | 0.006 |
| *Attention* | 0.011 | 0.0099 | 86 | 0.24 |

### RQ 3: Relation between physiological and behavioral engagement

From the results of RQ1 and RQ2, we can conclude that behavioral and physiological listening engagement are related. As a next step, we analyzed this in more depth and evaluated whether this relationship holds within each condition. Using NLME models, we tested whether synchronization predicts behavioral listening engagement while controlling for the target and distractor. Figure 6 shows inconsistent patterns across body modalities. Brain synchronization increases as subjective listening engagement increases across all conditions. The NLME revealed a significant relationship even after controlling for conditions. Second, for HR synchronization, the same pattern is observed in the non-engaging conditions but not in the engaging ones. No significant main effect of synchronization was identified, nor was there an interaction effect, contrary to our expectations based on the figures. The EDA NLME model revealed no main effect of interpersonal synchronization on behavioral engagement. However, there was a small but significant interaction effect between the distractor and EDA interpersonal synchronization, suggesting that the relationship between EDA and behavioral engagement was present only when the target was non-engaging. For the full model output, we refer to Table 7.

**Figure 6:**
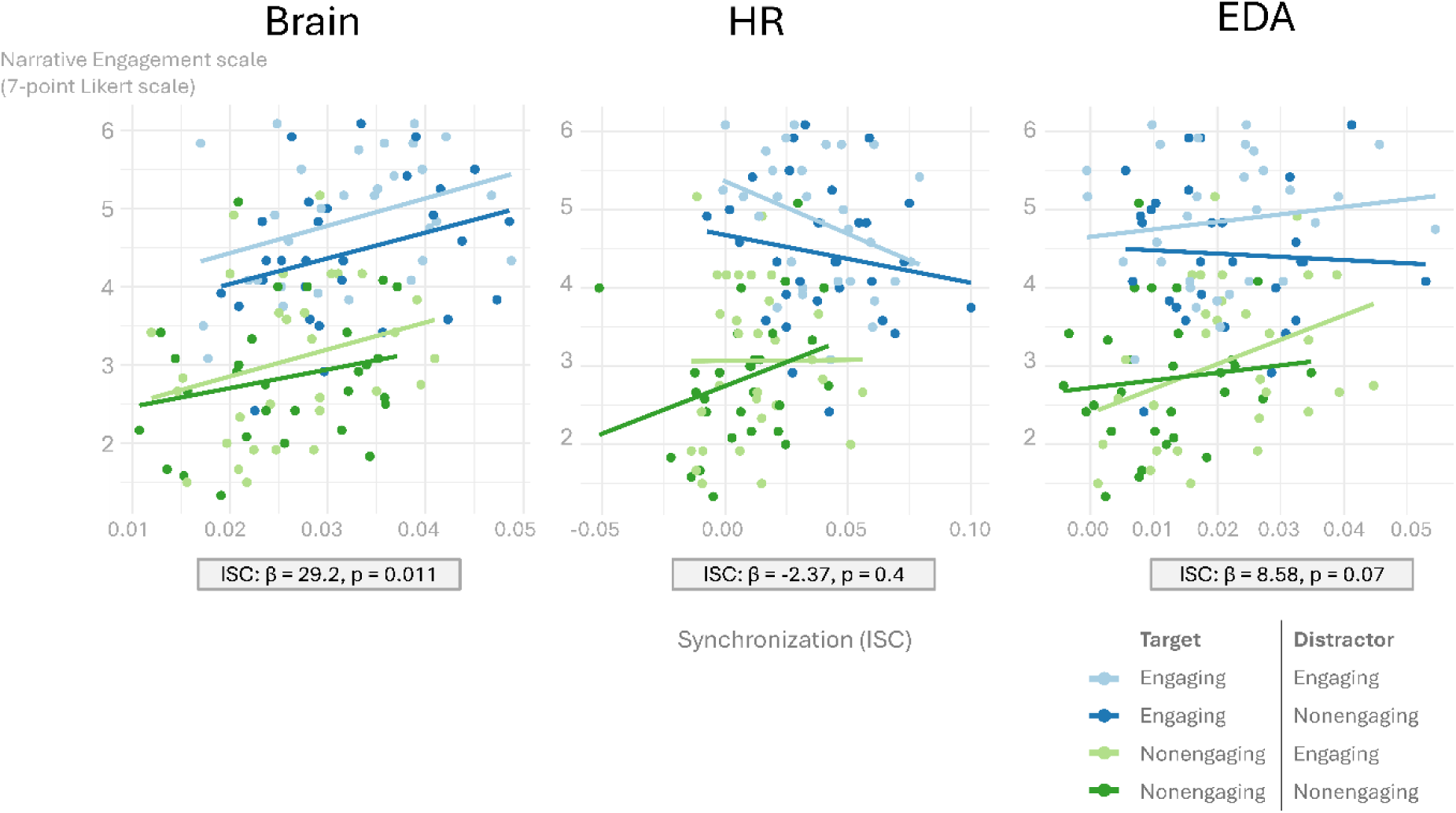
When controlled for the experimental conditions (i.e., induced listening engagement), the relationship between self-reported listening engagement and interpersonal synchronization remains, however, only for brain response and not for HR or EDA. This figure contains scatter plots showing the relationship between interpersonal synchronization and self-rated listening engagement for each condition individually.

**Table 7:** Non-linear mixed effect models for the behavioral – objective relationship.

| **Brain model** |  |  |  |  |
| --- | --- | --- | --- | --- |
| *Listening Engagement ~ EEG ISC + target + distractor + target \* synchronization + distractor \* synchronization + (1 \| participants)* |  |  |  |  |
|  | *Estimate* | *Std Err* | *df* | *p-value* |
| *Intercept* | 2.95 | 0.39 | 83 |  |
| *Synchronization* | 29.2 | 13 | 83 | 0.011 |
| *Target* | 0.75 | 0.23 | 83 | 0.005 |
| *Distractor* | 0.026 | 0.22 | 83 | 0.9 |
| *Synchronization\* Target* | 0.24 | 7.73 | 83 | 0.98 |
| *Synchronization\* Distractor* | 4.76 | 7.14 | 83 | 0.56 |
| **HR Model** |  |  |  |  |
| *Listening Engagement ~ HR ISC + target + distractor + target \* synchronization + distractor \* synchronization + (1 \| participants)* |  |  |  |  |
|  | *Estimate* | *Std Err* | *df* | *p-value* |
| *Intercept* | 3.9 | 0.16 | 83 |  |
| *Synchronization* | -2.37 | 3.57 | 83 | 0.4 |
| *Target* | 0.96 | 0.11 | 83 | 0.001 |
| *Distractor* | 0.19 | 0.087 | 83 | 0.085 |
| *Synchronization\* Target* | -3.22 | 3.44 | 83 | 0.5 |
| *Synchronization\* Distractor* | -1.02 | 2.54 | 83 | 0.73 |
| **EDA model** |  |  |  |  |
| *Listening Engagement ~ EDA ISC + target + distractor + target \* synchronization + distractor \* synchronization + (1 \| participants)* |  |  |  |  |
|  | *Estimate* | *Std Err* | *df* | *p-value* |
| *Intercept* | 3.62 | 0.17 | 83 |  |
| *Synchronization* | 8.58 | 6.12 | 83 | 0.07 |
| *Target* | 0.97 | 0.12 | 83 | 0.001 |
| *Distractor* | -0.094 | 0.12 | 83 | 0.49 |
| *Synchronization\* Target* | -6.1 | 5.61 | 83 | 0.55 |
| *Synchronization\* Distractor* | 12.1 | 5.72 | 83 | 0.04 |

## Discussion

Attention and listening engagement remain difficult to differentiate in the field of speech processing. In addition, interpersonal synchronization is an emerging neurophysiological measure that has been suggested to reflect both, making its interpretation challenging. With this study, we sought to unravel the relative contributions of attention and listening engagement to interpersonal synchronization (via ISC). We subjected participants to an auditory attention paradigm in which they listened simultaneously to two continuous speech stimuli, and we used AAD to measure attention. We manipulated listening engagement by using both engaging and non-engaging stimuli in the target and distractor stimuli, and included behavioral measures of listening engagement as a ground truth.

### Behavioral and physiological measures of listening engagement are sensitive to listening <u>engagement manipulations</u>

We manipulated listening engagement by selecting stimuli categorized as engaging or non-engaging based on factors known to influence engagement (e.g., content interest (Herrmann & Johnsrude, 2020; Lambrechts et al., 2025). Engaging stimuli consisted of segments from a popular Flemish podcast, whereas non-engaging stimuli were excerpts from parliamentary debates from 2015, making them no longer relevant. Ratings on the Narrative Engagement Scale (Busselle & Bilandzic, 2009) confirmed the stimuli’s effectiveness in inducing the desired engagement state, replicating earlier findings (Lambrechts et al., 2025). Notably, ignored speech also affected participants’ engagement ratings, despite explicit instructions to base them only on the target speech. This outcome suggests that irrelevant background input can influence perceived engagement, which is consistent with previous findings (Alamir & Hansen, 2021; Carpentier, 2010). In contrast, interpersonal synchronization was driven by target speech across all modalities (brain, HR and EDA). To a small extent, EDA synchronization was also driven by the distractor. From previous research, we know that attending to speech increases interpersonal synchronization, whereas not paying attention to speech decreases it (Madsen & Parra, 2022, 2025). This explains why, in this experiment, distractor speech did not influence brain and HR interpersonal synchronization, because it was not sufficiently processed in the absence of attention. The different pattern observed for EDA synchronization suggests potential modality-specific differences in interpersonal synchronization, though this should be interpreted cautiously given the small effect. These findings appear to diverge from the behavioral engagement ratings. Yet, it is possible that the behavioral rating is more sensitive to bias, induced by the distractor, whereas interpersonal synchronization is not. Generally, the observed synchronization was similar in magnitude to that in previous studies and showed significant synchronization in each of the modalities, especially for HR and brain interpersonal synchronization (Irsik et al., 2022; Lambrechts et al., 2025; Madsen & Parra, 2022). This supports the idea that ISC and, by extension, interpersonal synchronization are robust measures of listening engagement, even in the presence of an additional distracting speech stimulus.

### Listening engagement modulates attention

Across all conditions, we conclude that participants selectively attended to the required stimulus. This was reflected in AAD accuracies of around 0.7, with the majority being significant, which is in line with previous research (Geirnaert et al., 2021). From the AAD accuracies, we derived two things: 1) we found that attention was higher when the target was engaging, and 2) attention was higher when the distractor was non-engaging. This finding suggests that selective attention is influenced by both the speech we are motivated or instructed to listen to and the background sounds. Previous research showed that an uninteresting/non-engaging distracting stimulus is easier to ignore than an engaging one (Carpentier, 2010), so this finding was well within our expectations. If our stimulus manipulation had perfectly manipulated listening engagement but not attention, there would be no effect of engagement on AAD performance. However, as discussed in the introduction, attention and listening engagement are closely related, making it challenging to manipulate engagement without also affecting attention (Bradbury, 2023; Madsen & Parra, 2022). The differences in outcomes between engagement and attention results suggest that both measures reflect related yet distinct phenomena. In the following analyses, we aimed to further unravel this.

### Interpersonal synchronization as a measure of attention and listening engagement

The main aim of this study was to assess what interpersonal synchronization reflects: attention, listening engagement or a combination of both. Previous research investigating attention and listening engagement in isolation had already linked synchronization to both concepts (Lambrechts et al., 2025; Madsen & Parra, 2025). However, no formal comparison combining both manipulations within a single experiment has been conducted. If attention and engagement explain unique variance in synchronization, this would provide stronger evidence that they represent partially distinct constructs. We used validated measures of engagement (NES scores (Busselle & Bilandzic, 2009) and attention (AAD accuracy (Geirnaert et al., 2021)) as predictors. Listening engagement contributed significantly and uniquely to interpersonal synchronization across neural, HR and EDA signals, supporting its validity as a physiological index. This aligns with earlier findings, including our own (Lambrechts et al., 2025), linking behavioral and physiological engagement measures. Attention also uniquely predicted interpersonal synchronization; however, this was true only for neural synchronization, not for HR or EDA responses. This may indicate that cognitive aspects of engagement are primarily reflected in neural activity, whereas physiological measures such as HR and EDA are more sensitive to affective and arousal-related components (Anthiyur Aravindan et al., 2023; Kołodziej et al., 2019; Marois et al., 2024). Contrary to our results, however, HR synchronization has previously been associated with attention (i.e., lower interpersonal synchronization when people were distracted from listening, than when they paid attention (Pérez et al., 2021)). Consequently, it remains unclear what the HR responses actually reflect. In this context, it is important to note that manipulating attention likely entails a simultaneous manipulation of engagement, whereas the reverse is not necessarily true. This raises the possibility that effects previously attributed to attention may have actually reflected engagement (Irsik et al., 2022; Madsen & Parra, 2022). This hypothesis is supported by our previous publication, which suggested that HR synchronization increased with engagement but not with attention (Lambrechts et al., 2025).

Further research should delve deeper into the processes at play in this specific scenario. Overall, engagement showed statistically reliable effects across more analyses, although the absolute effect estimates were generally smaller. These findings collectively suggest that interpersonal synchronization reflects both listening engagement and attention, indicating that the two constructs are partially distinct yet overlapping, and may have additive effects across physiological modalities.

### Interpersonal synchronization as a measure of listening engagement across conditions

Interpersonal synchronization provided additional insight into listening engagement beyond the experimental condition. When modeling behavioral listening engagement (NES scores) while controlling for both target and distractor stimuli, brain synchronization remained a significant predictor of engagement. So far, to our knowledge, only a few studies have reported analyses of concordance between behavioral and physiological engagement; the relation between the two has not been proven (Czepiel et al., 2025; Irsik et al., 2022). In a previous report, we demonstrated this relationship by showing significant correlations between both engagement indices (Lambrechts et al., 2025). In the present study, we take this one step further by showing that this relationship holds even after controlling for the additional manipulations we made. This suggests that neural synchronization reflects an actual state of listening engagement and is not merely a stimulus-driven effect, supporting its potential as an objective measure of listening engagement. In contrast, EDA and HR synchronization did not show clear or significant relationships with behavioral engagement after controlling for the experimental manipulations of attention and overall engagement of the speech material. Data visualization suggested a possible interaction effect, whereby the association with engagement appeared negative when the attended stimulus was engaging and positive when it was not. This interaction was significant for EDA synchronization. Despite the lack of a statistically significant effect for HR synchronization, these results raise interesting questions about stimulus-dependent dynamics that require further investigation. Several factors may explain the weaker findings for HR and EDA. These measures may be less sensitive to engagement than brain synchronization. Additionally, limited data per condition and noise introduced by presenting two stimuli simultaneously, may have reduced the reliability and stability of interpersonal synchronization. Nonetheless, these findings extend our previous work by suggesting that measures of listening engagement are influenced not only by the experimental manipulations themselves but also by how those manipulations influence the listener’s subjective experience.

### Limitations

Despite repeated instructions to maintain attention to the target stimulus, we cannot guarantee that all participants consistently complied with these instructions throughout the experiment. To encourage attentiveness, several precautions were implemented, including the randomization of conditions, offering breaks as needed, and informing participants that a brief content-retention questionnaire would follow each listening condition to maintain motivation. Additionally, based on our literature review, we recognize that listening engagement and, by extension, interpersonal synchronization may not be adequately captured as a purely categorical construct, particularly in applied contexts. Creating stimuli that differ only in listening engagement is inherently challenging, as engagement is influenced by multiple interacting factors. To partially control for this, we selected recordings featuring only female speakers in natural speaking contexts, rather than self-produced, artificial material. Nevertheless, categorizing stimuli into ‘high’ and ‘low’ engagement conditions may still appear somewhat simplified or artificial. Despite this limitation, the present study represents an important step toward developing more continuous and ecologically valid measures of listening engagement.

## Conclusion

The goal of this paper was to better understand exactly what interpersonal synchronization reflects in an auditory context and to provide evidence that attention, as well as listening engagement, can explain unique variance found in this measure. We selected our stimuli to be either engaging or non-engaging and validated this manipulation using a behavioral measure (narrative engagement scale). We found that both attention and listening engagement contribute unique variance to neural interpersonal synchronization during listening, confirming our hypothesis. This suggests that they are distinct constructs that create complementary listening experiences. For interpersonal synchronization of EDA and HR responses, we found it was significantly driven by listening engagement but not by attention. These findings suggest that brain interpersonal synchronization reflects attention and the cognitive component of listening engagement, whereas body interpersonal synchronization reflects only the affective component of listening engagement. Additionally, our analysis of the relationship between behavioral listening engagement and interpersonal synchronization supports the use of neural IS as an objective measure of listening engagement, independently of any experimental manipulation.

### Artificial Intelligence Statement

Grammarly (*Grammarly Writing Assistant*) and ChatGPT (*ChatGPT*, 2026) were used as writing support tools to assist with correcting spelling and grammatical errors and to enhance the clarity and fluency of the text. The resulting text was revised by the authors prior to inclusion in the manuscript.

## Acknowledgments

The authors thank Mirne Vandenbruaene and Marlies Vandermeulen for their help during data collection.

## Data availability statement

The datasets produced during the current study are not publicly available due to the personal nature of the data, but are available from the corresponding author on reasonable request.

## Author contributions

**L.L.:** Conceptualization, Methodology, Software, Data curation, Formal analysis, Investigation, Validation, Visualization, Writing – original draft, Writing – review and editing. **B.A.:** Conceptualization, Methodology, Data curation, Investigation, Writing – review and editing, Supervision. **J.V.** Conceptualization, Methodology, Formal analysis, Writing – review and editing, Supervision, Project Administration. **B.B.** Conceptualization, Writing – review and editing, Supervision. **T.F.** Conceptualization, Resources, Writing – review and editing, Supervision, Project Administration, Funding acquisition.

